# Estimation of Global and Local Complexities of Brain Networks: A Random Walks Approach

**DOI:** 10.1101/733725

**Authors:** Roberto C. Sotero, Lazaro M. Sanchez-Rodriguez, Narges Moradi

## Abstract

The complexity of brain activity has been observed at many spatial scales and there exists increasing evidence supporting its use in differentiating between mental states and disorders. Here we proposed a new measure of network (global) complexity that is constructed as the sum of the complexities of its nodes (i.e, local complexity). The local complexity of each node is regarded as an index that compares the sample entropy of the time series generated by the movement of a random walker on the network resulting from removing the node and its connections, with the sample entropy of the time series obtained from a regular lattice (the ordered state) and an Erdös-Renyi network (disordered state). We studied the complexity of fMRI-based resting-state functional networks. We found that positively correlated, or “**pos**”, network comprising only the positive functional connections has higher complexity than the anticorrelation (“**neg”**) network (comprising the negative functional connections) and the network consisting of the absolute value of all connections (“**abs”**). We also found a significant correlation between complexity and the strength of functional connectivity. For the **pos** network this correlation is significantly weaker at the local scale compared to the global scale, whereas for the **neg** network the link is stronger at the local scale than at the global scale, but still weaker than for the **pos** network. Our results suggest that the **pos** network is related to the information processing in the brain and should be used for functional connectivity analysis instead of the **abs** network as is usually done.

## 1. Introduction

The development of a quantitative measure of complexity has proven difficult because of the variety of systems that may be labelled as ‘complex’. In the case of the complexity of networks, perhaps the most popular approach has been the use of information-based measures (Bonchev & Buck, 2005; Dehmer, Barbarini, Varmuza, & Graber, 2009). The basic principle to construct these measures is to select an arbitrary graph invariant *X*, partitioned as *x*_1_, …, *x*_*N*_. Probabilities can be inferred for each partition using the entities 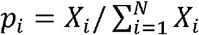 since it holds that 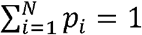. The information content of the graph is then computed using the Shannon formula (Shannon, 1948): 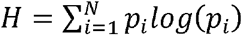. Another important definition of complexity was proposed by Kolmogorov (Kolmogorov, 1968). The Kolmogorov complexity of a network is the length of the shortest computer program that produces the network as output. Although Kolmogorov complexity is uncomputable it can be approximated to a degree that allows its practical use (Li & Vitányi, 2008).

The measures of complexity described above assume it to be a monotonically increasing function of disorder. However, complexity can also be defined as a monotonically increasing function of order, as shown by McShea (McShea, 1991), who found that the morphological complexity of organisms changed with the level of self-organization, and the latter with order. Finally, complexity can be defined as a convex function of disorder; i.e., a quantity that attains a minimum for both completely ordered and completely disordered systems, and a maximum at some intermediate level of disorder or order (López-Ruiz, Mancini, & Calbet, 1995; Shiner, Davison, & Landsberg, 1999; Tononi, Edelman, & Sporns, 1998). Here, we adopt this latter notion by assuming that network complexity achieves a minimum for random networks, also known as Erdös-Renyi (ER) networks (Erdös & Rényi, 1959), and regular lattice (RL) networks (Watts & Strogatz, 1998).

In addition to the global complexity of the brain network, in this work we are interested in computing the local complexities (a measure for each of the different brain areas), such that the global complexity of the network is the sum of the local ones; i.e, the complexity of the system is the sum of the complexity of its parts. To estimate the complexities, we let a random walker diffuse on the network and construct a time series of the strengths of the nodes (brain areas) visited by the walker. The sample entropy (SampEn) (Richman & Moorman, 2000) of the time series is then calculated. Local complexities are obtained by iteratively removing a node and all its connections, constructing the time series from the walker movement in the resulting network, computing the SampEn, and comparing this value to the average value obtained from 1000 ER and 1000 RL networks with the same degree distribution and connections strengths.

Functional connectivity in the brain is defined as the synchronization of neurophysiological events among anatomically separated brain areas (Friston, Jezzard, & Turner, 1994). Biswal and colleagues (Biswal, Yetkin, Haughton, & Hyde, 1995) were the first to report that during resting-state the primary motor regions in the left and right hemispheres were positively correlated. Later studies identified positive correlations between regions that are now known to comprise the default mode network (DMN) (Buckner, Andrews-Hanna, & Schacter, 2008; Raichle et al., 2001). In addition to the reported correlated networks, anticorrelated networks have also been reported by several studies (Michael D. Fox, Zhang, Snyder, & Raichle, 2009; Gopinath, Krishnamurthy, Cabanban, & Crosson, 2015; Liang, King, & Zhang, 2012). Although anticorrelations have been attributed to the global signals removal recent studies suggest a physiological basis (Michael D. Fox et al., 2009; Kazeminejad & Sotero, 2019). For this reason, in this paper we computed three different functional connectivity matrices for each subject using the Pearson correlation between the resting-state functional magnetic resonance imaging (fMRI) signals recorded from each of the 116 brain areas considered. A matrix consisting of the absolute value of all connections (denoted as **abs**), a matrix consisting of only the positive connections (denoted as **pos**) representing the positively correlated network, and a matrix comprising the absolute value of only the negative connections (denoted as **neg**) representing the anticorrelation network. We then compute the local complexities of the 116 brain areas, as well as the global complexities of the entire brain network, and seven known functional networks of the brain (Sedeño et al., 2016): default mode network (DMN), frontoparietal (FP), salience (SAL), sensorimotor (SM), visual (V), cerebellar (CER), and temporo-basal-ganglial (TBG) networks. Our results show that the **pos** network has higher global complexity than the **neg** and **abs** networks. We also found a link between complexity and functional connectivity which changes with the spatial scale and the type of brain network. For the **pos** network this link is significantly weaker at the local scale compared to the global scale, whereas for the **neg** network the link is stronger at the local scale than at the global scale, but still weaker than for the **pos** network. Also in the **pos** network global complexity was strongly correlated to the network integration and segregation, whereas **neg** was not significant correlated with integration and segregation. The network formed by taking the absolute value of the functional connectivity (**abs**) presented lower correlations than the **pos** case. Our results suggest that the **pos** network is related to the information processing in the brain network and should be used for functional connectivity analysis instead of the **abs** network.

## 2. Methods

### 2.1. Data acquisition and preprocessing

The resting-state fMRI dataset of 89 subjects from the NIH Human Connectome Project (HCP) (https://https://db.humanconnectome.org) (Van Essen et al., 2013) is used in this research. Each subject was involved in 4 runs of 15 minutes each using a 3 T Siemens scanner, while their eyes were open and had a relaxed fixation on a projected bright cross-hair on a dark background. The data were acquired with 2.0 mm isotropic voxels for 72 slices, TR=0.72 s, TE=33.1 ms, 1200 frames per run, 0.58 ms echo spacing, and 2290 Hz/Px bandwidth (Moeller et al., 2010). Therefore, the fMRI data were acquired with a spatial resolution of 2 × 2 × 2 mm and a temporal resolution of 0.72 s, using multibands accelerated echo-planar imaging to generate a high quality and the most robust fMRI data. The fMRI data were spatially preprocessed to remove spatial artifacts produced by head motion, B_0_ distortions, and gradient nonlinearities (Jovicich et al., 2006). Since comparison of fMRI images across subjects and studies is possible when the images have been transformed from the subject’s native volume space to the MNI space, fMRI images were wrapped and aligned into the MNI space with FSL’s FLIRT 12 DOF affine and then a FNIRT nonlinear registration (Jenkinson, Bannister, Brady, & Smith, 2002) was performed. In this study, the MNI-152-2 mm atlas (Mazziotta et al., 2001) was utilized for fMRI data registration.

### 2.2. Construction of functional connectivity matrices

The peak voxel in each region, that is, the voxel of maximal activation, was selected by computing the Root Mean Square (RMS) for each voxel's fMRI signal over all time. It has been shown that the peak voxel provides the best effect of any voxel in the ROI (Sharot, Delgado, & Phelps, 2004). Additionally, the peak voxel activity correlates well with evoked scalp electrical potentials than approaches that average activity across the ROI. This means that the peak voxel represents the ROI’s activity better than other choices (Arthurs & Boniface, 2003). The peak voxel in each region is determined using previously published Talairach coordinates (after conversion to MNI coordinates and using AAL 116 atlas) (M. D. Fox et al., 2005). The resulting signal was filtered to keep only low frequency fluctuations (0.01–0.08 Hz) (Yan & Zang, 2010). Finally, the global signal (i.e., the average of the fMRI signals over the whole brain (Michael D. Fox et al., 2009)) was regressed out.

We then computed the Pearson correlation between all possible pairs of time series creating a 116×116 functional connectivity matrix for each subject. Three different networks were obtained from this matrix. A network consisting of the absolute value of all connections (denoted as **abs**) which is the most commonly used in fMRI connectivity studies (Meier et al., 2016; Meszlényi, Hermann, Buza, Gál, & Vidnyánszky, 2017; Salvador et al., 2005), a network consisting of only the positive connections (denoted as **pos**), and a network comprising the absolute value of only the negative connections (denoted as **neg**). In all cases *p*-values were corrected by means of a multiple comparison analysis based on the false discovery rate (FDR) (Benjamini & Hochberg, 1995).

### 2.3. Construction of the time series of the random walker’s movements on the connectivity matrix

We first consider an unweighted network consisting of *N* nodes. We place a large number *K*(*K* ≫ *N*) of random walkers onto this network. At each time step, the walkers move randomly (with the same probability) between the nodes that are directly linked to each other. We allow the walkers to perform *T* time steps. As a walker visits a node, we record the degree of the node. Thus, after *T* time steps, we obtain *K* time series reflecting different realizations of the random walker’s movement on the network. Nodes with high degree (hubs) will appear more frequently in the series than nodes with low degree.

In the case of weighted networks, such as the functional connectivity matrix representing the brain network, the transition probability *P*_*ij*_ from brain area *i* to brain area *j* is given by 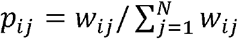, where *w*_*ij*_ is the weight of the connection from area *i* to area *j* (Zhang, Shan, & Chen, 2013). We then construct a time series with the strengths of the nodes *i* visited by the walker: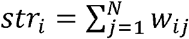.

### 2.4. Computing the entropy of the time series

In this paper we use sample entropy (SampEn) (Richman & Moorman, 2000) to estimate the complexity of the time series of the diffusion of the random walker in the network. SampEn improved from approximate entropy (ApEn) (Pincus, 1991) by reducing the bias caused by self-matching. For a time series *x*(*i*), 1≤ *i* ≤ *N*, of finite length *N*, we first reconstitute the *N* − *m* + 1 vectors *x*_*m*_(*i*) following the form:

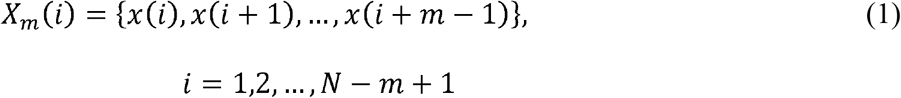

where *m* is the embedding dimension.

Let 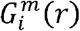 be the probability that any vector *x*_*m*_(*j*) is within distance *r* of *X*_*m*_(*i*):

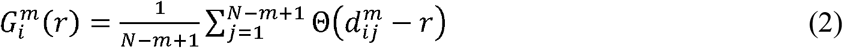

where 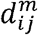 is the distance between the vectors *x*_*m*_(*i*) and *x*_*m*_(*j*), defined as:

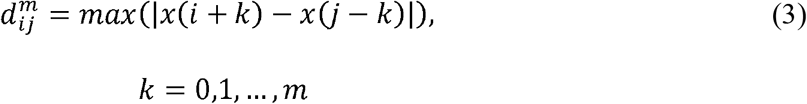

When the embedding dimension is *m*, the total number of template matches is:

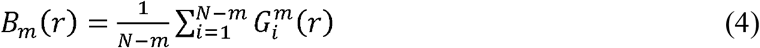

Similarly, when the embedding dimension is *m+1*, the total number of template matches is:

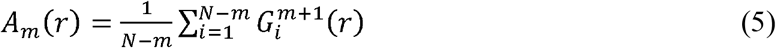

Finally, the SampEn of the time series is estimated by:

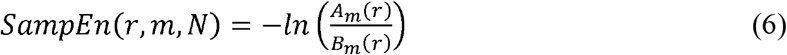

### 2.5. Computing local complexities and global complexity

We propose to obtain local complexities *c*_*i*_ by 1) iteratively removing a node and all its connections, 2) constructing the time series from the random walker diffusion in the resulting network, and 3) computing the SampEn of the time series obtained in the previous step. For node *i*, the resulting SampEn is labelled as *H*_≠i_. Then we compare this value to the average SampEn (computed followed the same procedure outlined before) of 1000 ER 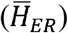 and 1000 RL networks 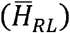 of the same size (i.e, *N*−1) and connections strengths taken from the original matrix. The local complexity is the percent this value is of the square of the entropy of the original matrix (*H*), multiplied by the probability (*p*_*i*_) of the appearance of the node in the time series:

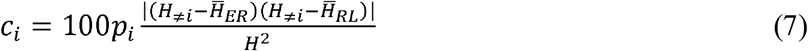

Figure 1 shows the steps described above for computing the local complexities. The global complexity of the network *C* is then computed as the sum of the local complexities:

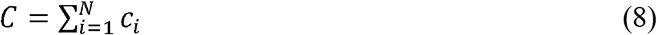

**Figure 1.**
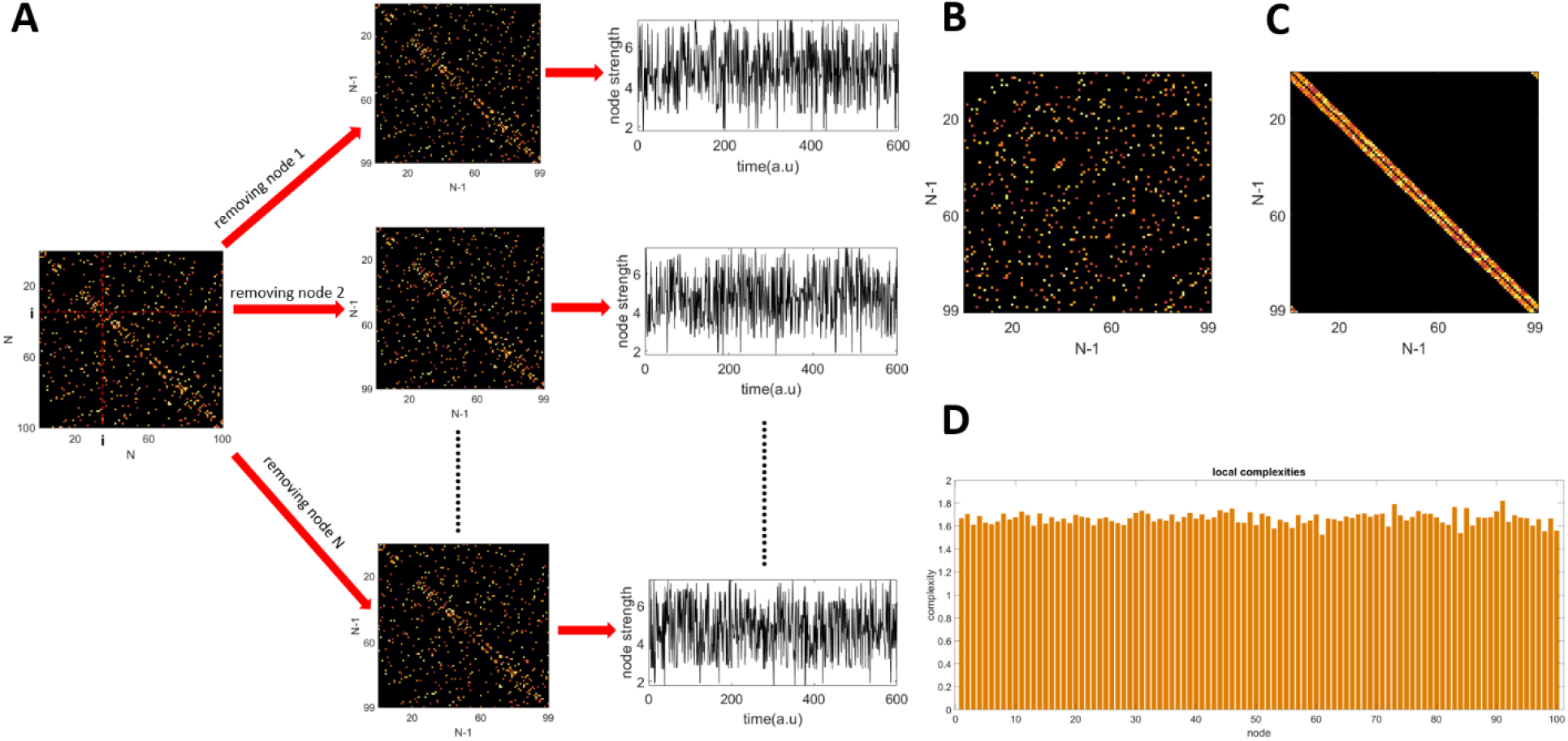
Methodology for computing local complexities. A) Given a connectivity matrix of size *N*, each node is removed iteratively and a new matrix of size (*N*−1)×(*N*−1) is obtained. Then a time series of node strengths is constructed from the diffusion of a random walker in the new matrix. B) A random network of size (*N*−1)×(*N*−1) with the same average degree and strengths as the matrices obtained in A. C) A regular network of size (*N*−1)×(*N*−1) with the same average degree and strengths as the matrices obtained in A. D) Local complexities.

## 3. Results

### 3.1.1 Global complexity of simulated complex networks

As stated before, the goal of this work is to propose a new measure of structural complexity that is useful for brain networks. To demonstrate the usefulness of the quantity we defined, we start by measuring how changes in the underlying network structure affects the observed values of global complexity. To this end, we devised a scenario in which the network gradually transforms from the perfectly orderly state (regular lattice network) to a completely random state (Erdos-Renyi network). Following equations (7) and (8) we expect complexity to have a minimum at these states. Network states different from these minimums would have a mixture of order and disorder and thus were modeled using the small-world model (Watts & Strogatz, 1998). In this model, nodes of the network are placed on a regular *k*-dimensional grid and each node is connected to *m* of its nearest neighbours, producing a regular lattice of nodes with equal degrees. Then, with probability *p*, each connection is randomly randomly rewired. The RL network corresponds to the value *p* = 0. When *p* > 0, edge rewiring is applied, and this changes the degree distribution of nodes. On the other end of the spectrum is the ER model (Erdös & Rényi, 1959), obtained when *p* = 1, in which there is no connectivity pattern between nodes. In between, SW networks, obtained for values 0 < *p* < 1, present high clustering and short path length (Watts & Strogatz, 1998).

Graph theoretical studies of mammalian cortical networks recreated from tract tracing experiments demonstrated that the cat and macaque interareal anatomical networks share similar small world properties of short path length and high clustering (Hilgetag & Kaiser, 2004; Sporns & Zwi, 2004). Additionally, studies of anatomical and functional connectivity networks estimated from human neuroimaging data also found small world characteristics (Bassett & Bullmore, 2006; Salvador et al., 2005). To simulate RL, SW and ER networks we use Matlab function ‘WattsStrogatz.m’ which has as inputs the parameters *k* and *p*.

Figure 2A shows examples of matrices of size *N* = 100, for five different values of the rewiring probability *P*, and three values of the mean node degree *k*. The weights in the network were generated from a uniform random distribution with values between 0 and 1. We then placed 10^4^ random walkers onto these networks. The steps for estimating the global complexity of the network are presented in Figure 1 and described in detail in the Methods section. Figure 2B shows the global complexity of a network as a function of the rewiring probability *p*. Three different values of the average node degree were used *k* = 6,8,10. The results show that for a fixed network size the maximum global complexity decreases with the increase of *k* (the network gets denser). Additionally, the probability at which the peak in complexity was achieved, also decreased with the increase of *k*.

**Figure 2.**
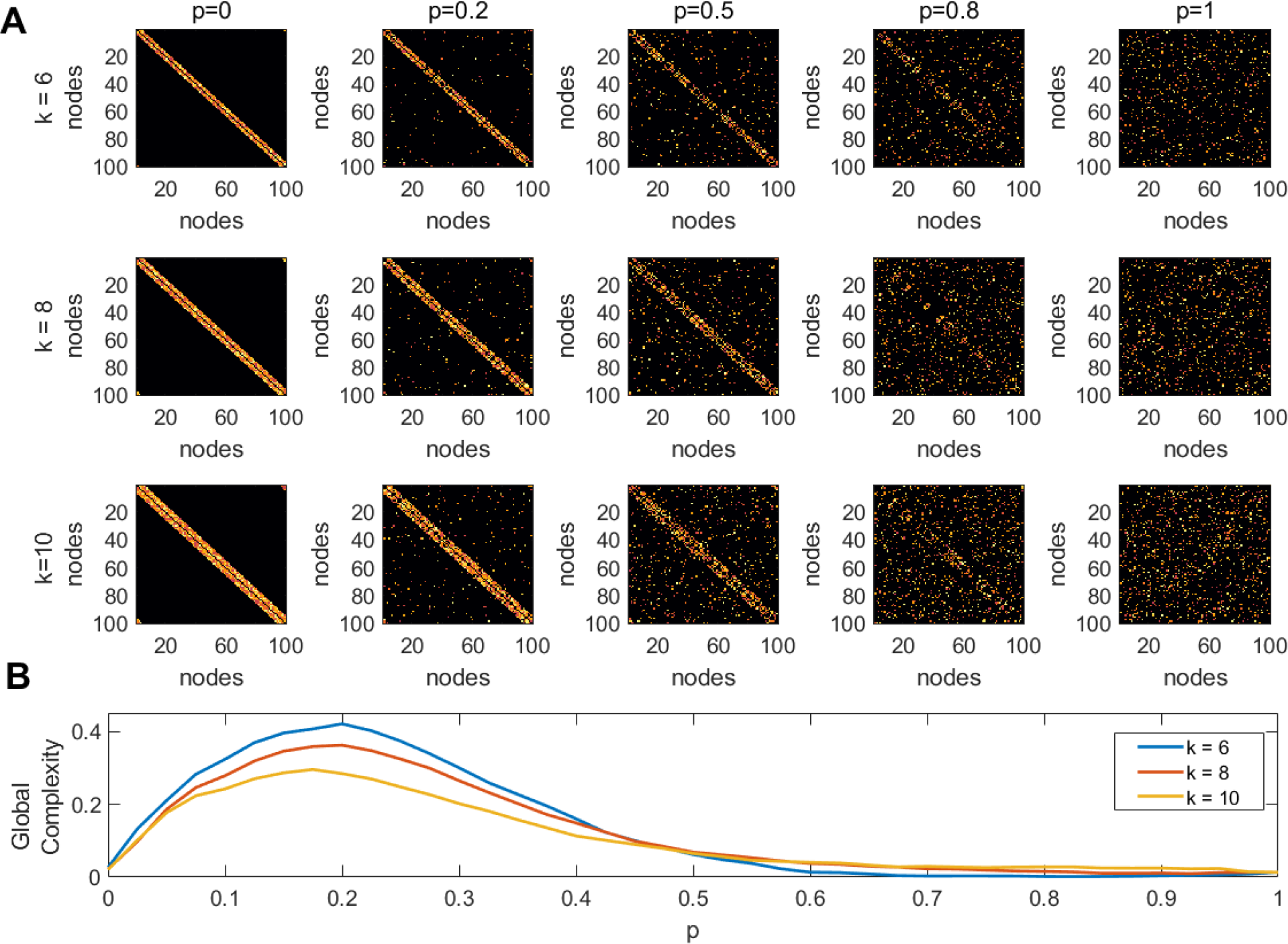
Global complexity of simulated networks. A) All networks have the same size N = 100, and were simulated using the Watts and Strogatz algorithm for creating small-world networks. The inputs to the model are the rewiring probability *P*, and mean node degree *k*. B) Global complexity as function of the rewiring probability *p*.

### 3.1.2 Complexity analysis of large-scale human brain networks

Figure 3A displays these matrices for one subject. Figure 3B shows the node degree of the three matrices average across all subjects, figure 3C shows their entropy, and figure 3D their global complexity. Our results show that the **pos** matrices are sparser than the **neg** matrices but have approximately the same entropy. This results in the **pos** network having a higher global complexity than the **neg** matrices. The **abs** matrices presented the lowest global complexity of the three cases.

**Figure 3.**
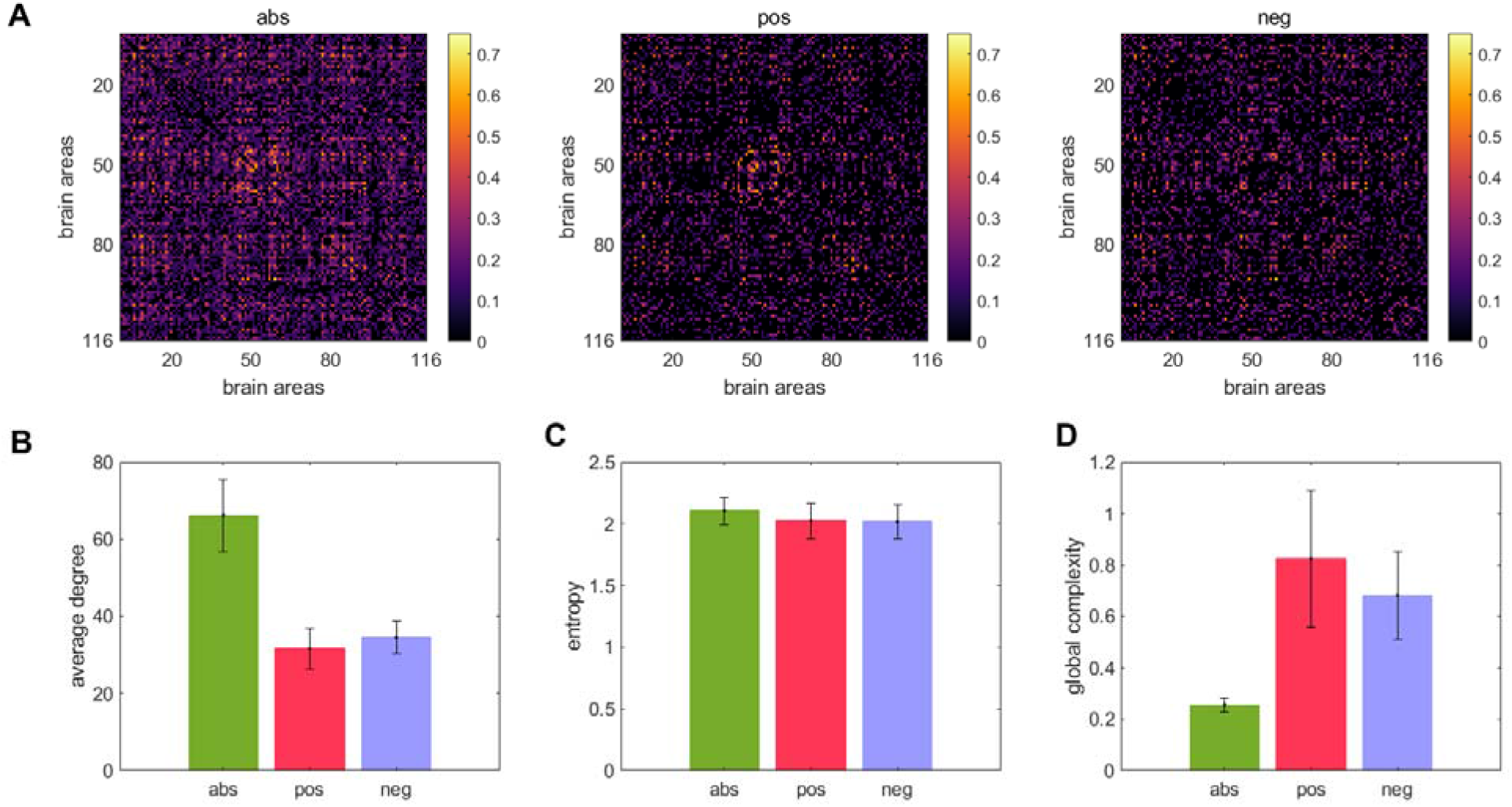
Global complexity of the entire brain network. A) A matrix consisting of the absolute value of all connections (denoted as **abs**), a matrix consisting of only the positive connections (denoted as **pos**), and a matrix comprising the absolute value of only the negative connections (denoted as **neg**). B) node degree averaged across subjects C) entropy averaged across subjects. D) global complexity averaged across subjects.

Figure 4A shows the linear fits between the global complexity and the sum of the functional connectivity strengths (SFCS) of the entire brain network for the **abs**, **pos**, and **neg** cases. We found that for the **pos** case, there is a strong correlation (*r* = 0.62, *p* = 9.5e^−11^) between global complexity and SFCS, followed by the **abs** case (*r* = 0.28, *p* = 0.0077). The anticorrelation network was not significantly correlated with SFCS (*r* = 0.11, *p* = 0.29). We also computed the linear fits between local complexities and the SFCS of each brain area (figure 4B). We found that for the **pos** case the link between complexity and functional connectivity was significantly weaker at the local scale compared to the global scale cr = 0.22, P = 2e^−23^). For the anticorrelation network the link was stronger at the local cr = 0.18, P = 3e^−21^) than at the global scale, but still weaker than for the **pos** case. The correlation between complexity and connectivity was essentially the same at the global and local scales for the **abs** case (figure 4).

**Figure 4.**
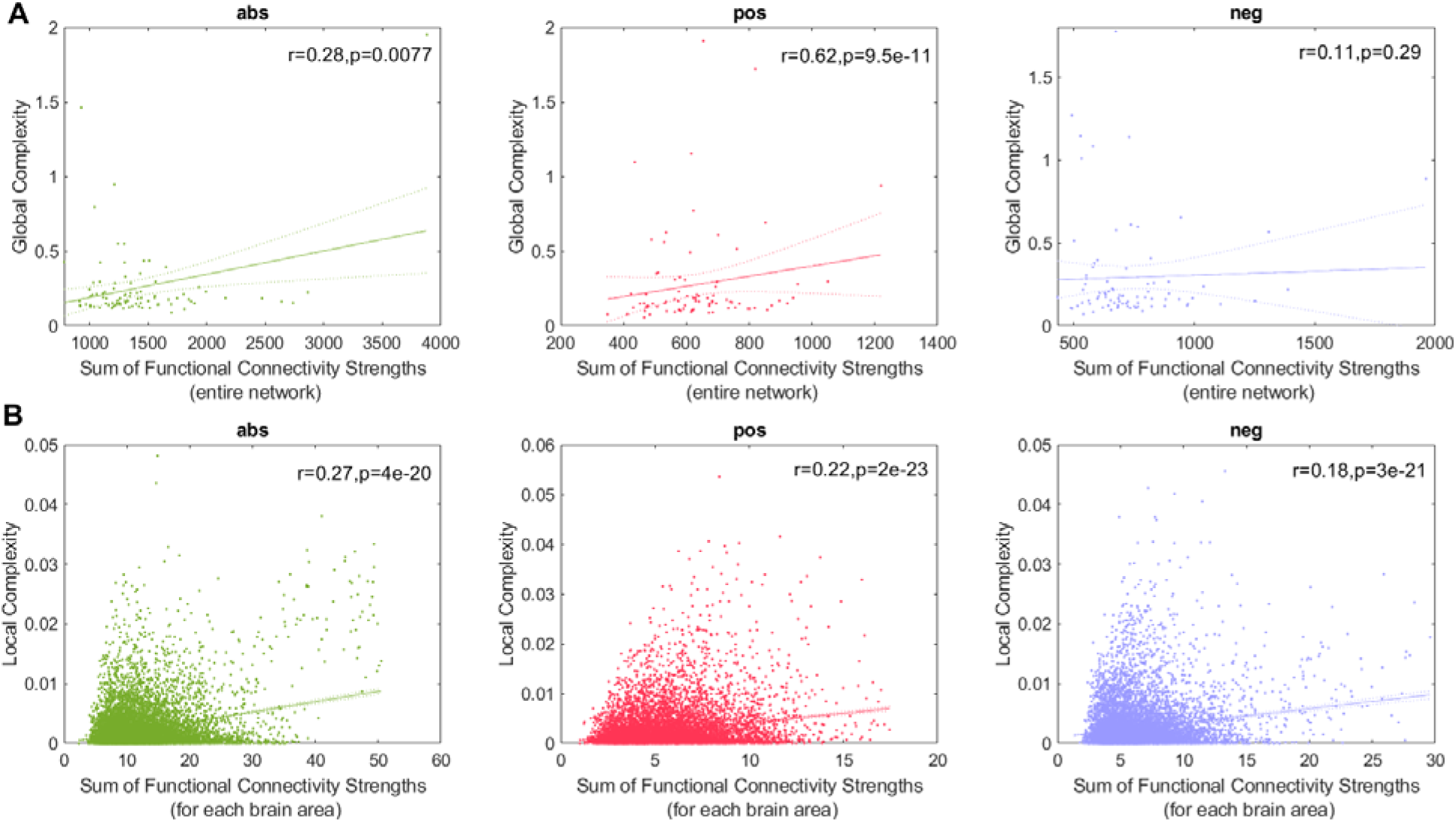
Link between Global complexity (A) and local complexities (B) and the sum of functional connectivity strengths. The **abs**, **pos** and **neg** networks appear in that order from left to right.

Complex networks are expected to present high values of both integration and segregation. Thus, we also explored the link between them and global complexity (figure 5). Integration and segregation were estimated using the global efficiency and average clustering coefficient of the network, respectively (Sporns, 2013). We found strong correlations between global complexity and both integration (*r* = 0.55, *p* = 2. 7e^−8^) and segregation (*r* = 0.59, *p* = 2e^−9^) for the **pos** network, and lower values for the **abs** case (r = 0.26, P = 0.016 for the correlation with integration and *r* = 0.28, *p* = 0.008 for the correlation with segregation). No significant correlations were found for the **neg** case.

**Figure 5.**
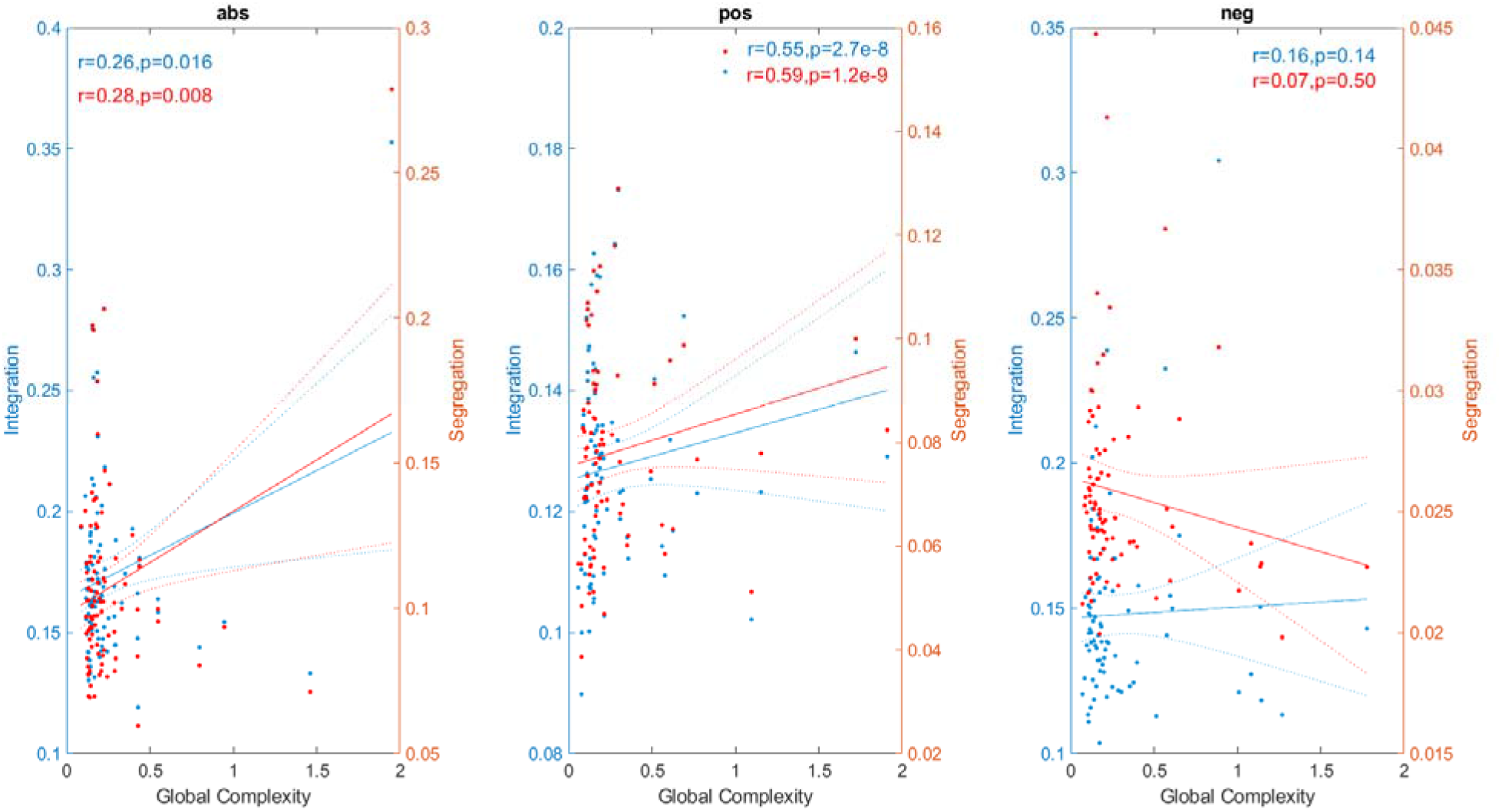
Link between global complexity and integration (blue) and segregation (red).

We also investigated the link between the three network types at the global (figure 6A) and local scales (figure 6B) finding that the **pos** and **neg** networks are not significantly correlated at any spatial scale.

**Figure 6.**
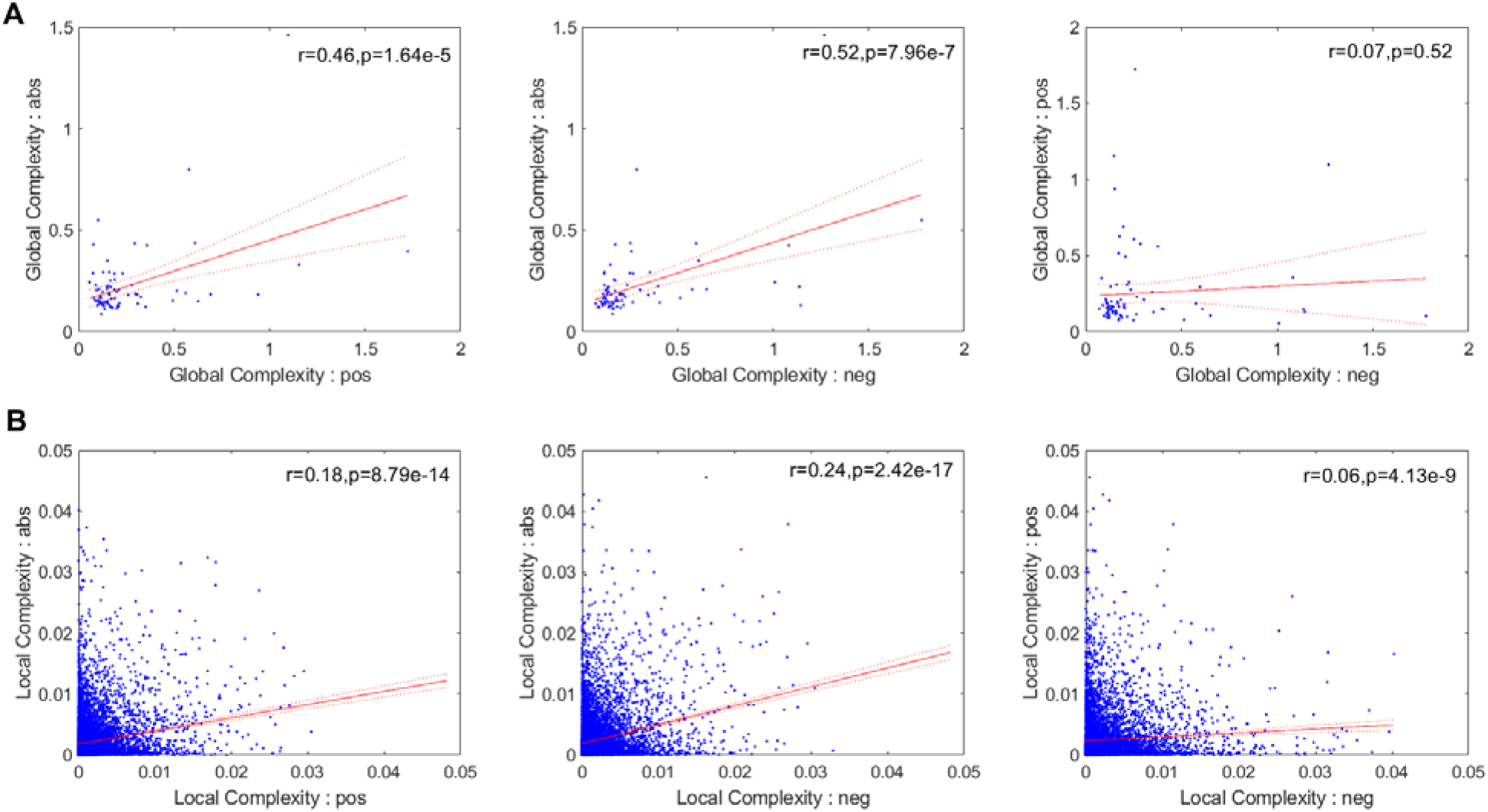
Link between the three network types (**abs**, **pos**, and **neg**) at the global (A) and local (B) scales.

Figure 7 presents the local complexity of the 116 brain areas for the **pos** and **neg** cases. Seven resting-state networks (Sedeño et al., 2016) were considered (DMN, FP, SAL, CER, V, SM, TGB) as well as areas that were not allocated to a network (NA). In both **pos** and **neg** cases, the area with the highest complexity belongs to the DMN (Angular R and Frontal Med Orb R). Figure 8 displays the local complexity for the **abs** case and the sum of the complexities of the **neg** and **pos** case (**neg+pos**). In the abs case, the highest complexity was obtained for the left paracentral lobule, while the right angular gyrus presented the highest complexity for the **neg+pos** case.

**Figure 7.**
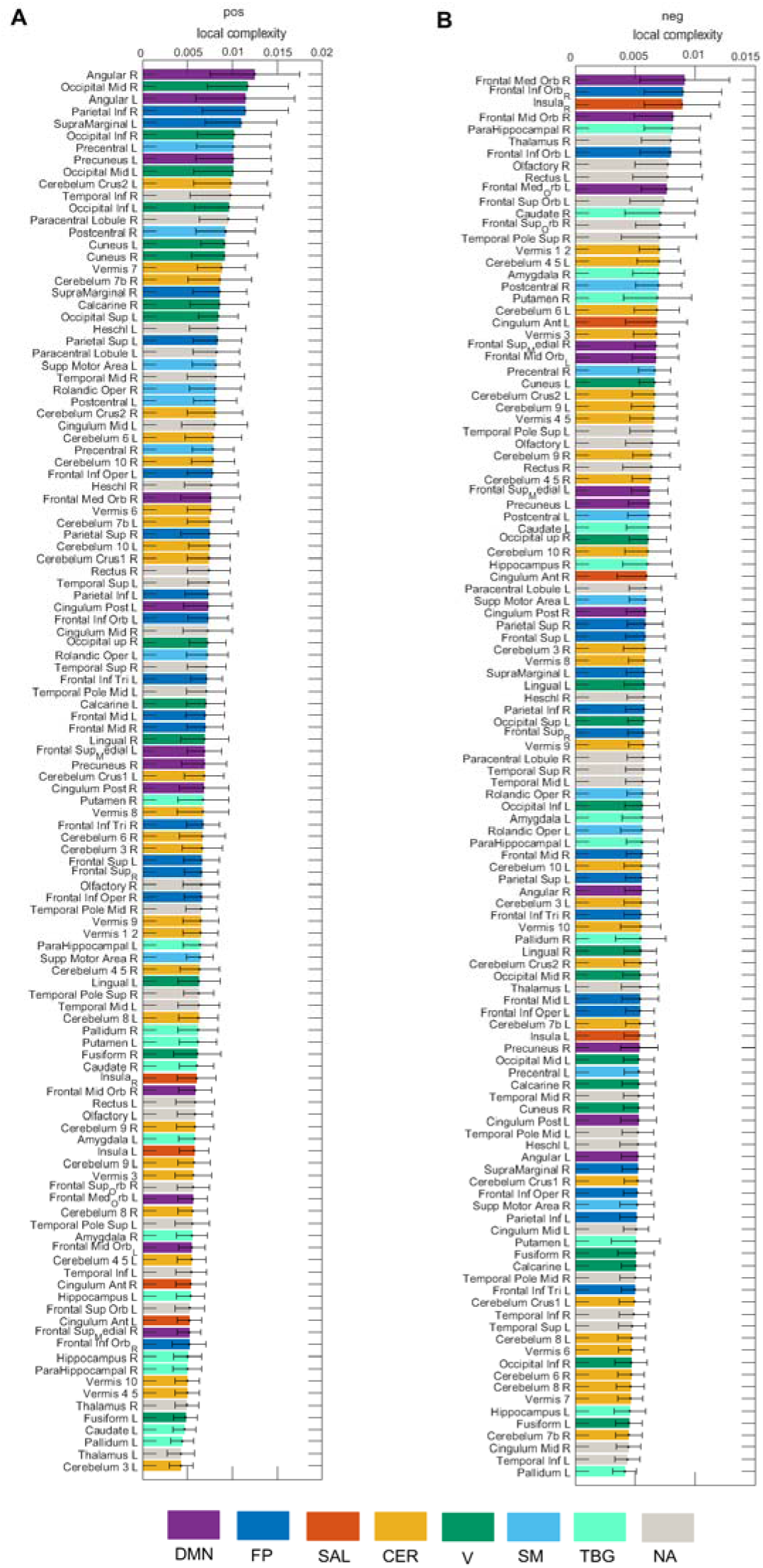
Local complexity of the 116 brain areas for the **pos** and **neg** cases. Seven resting-state networks (see Supplementary Table 1) are represented through different colors: default mode network (DMN), frontoparietal (FP), salience (SAL), sensorimotor (SM), visual (V), cerebellar (CER), and temporo-basal-ganglial (TBG) networks. The gray color represents areas not assigned (NA) to any of these networks.

**Figure 8.**
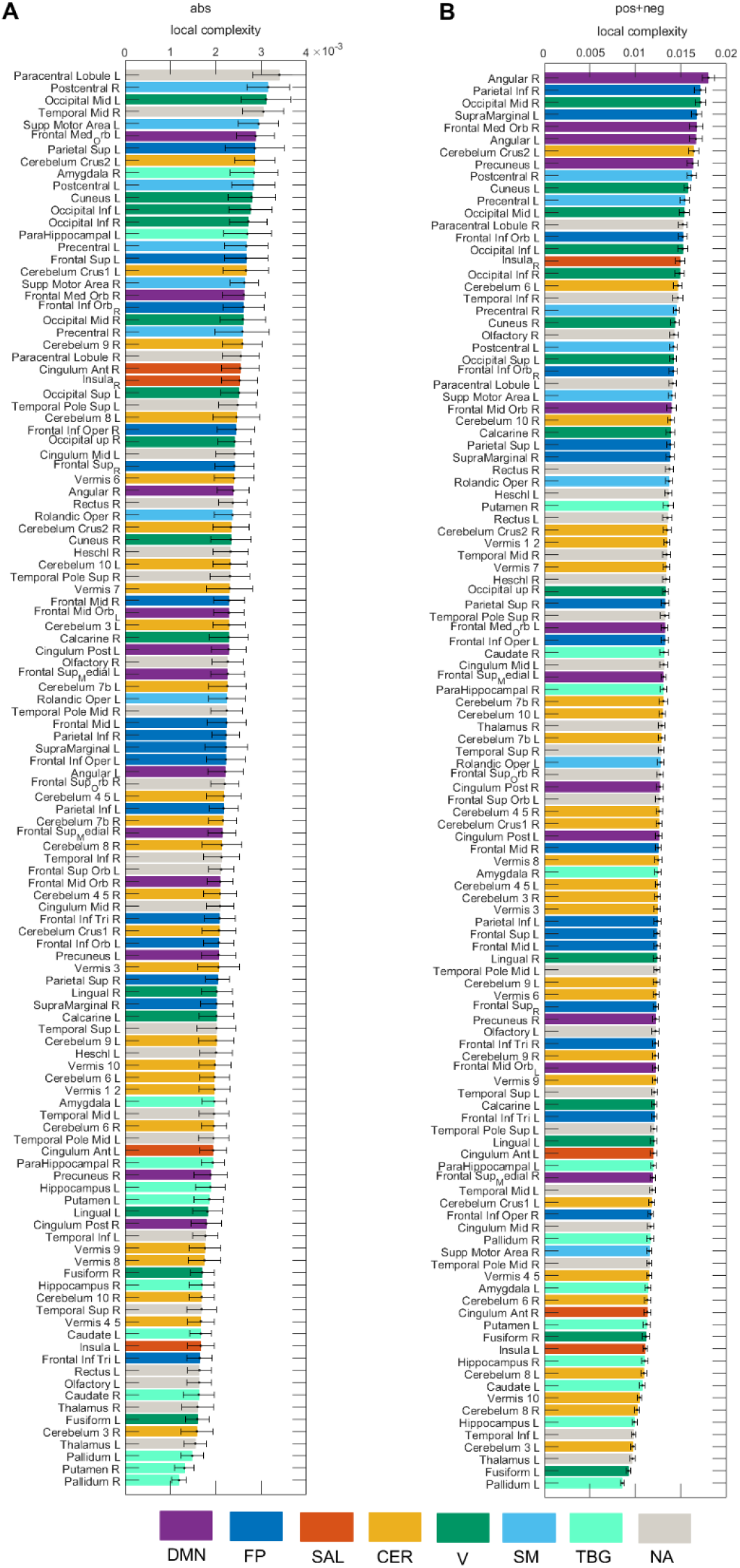
Local complexity of the 116 brain areas for the **abs** and **pos+neg** cases. Seven resting-state networks (see Supplementary Table 1) are represented through different colors: default mode network (DMN), frontoparietal (FP), salience (SAL), sensorimotor (SM), visual (V), cerebellar (CER), and temporo-basal-ganglial (TBG) networks. The gray color represents area not assigned (NA) to any of these networks.

We computed the global complexity of the seven resting-state networks (figure 9A). We found that the network with the highest complexity for all cases was the cerebellar network, while the network with the lowest complexity was the salience network. The DMN, FP, CER, V and SM networks presented more complexity in the **pos** than in the **neg** case, while the SAL and TGB networks were more complex in the **neg** case. When interpreting this result we need to be aware of the fact that since the global complexity of the network is computed as the sum of the local complexities (equation 8), networks comprising few brain areas (as is the case of the salience network) will have a low value of global complexity provided that the difference in the values of the local complexities is not high (see figures 7 and 8). To account for this issue, we also divided the global complexity of each network by the number of areas in each network (figure 9B). As a result, although the average contribution of the areas in the salience network to the network complexity is still the lowest among the seven resting state networks for the **pos** case, it is the areas in the visual network the ones with the lowest contribution in the **neg** case.

**Figure 9.**
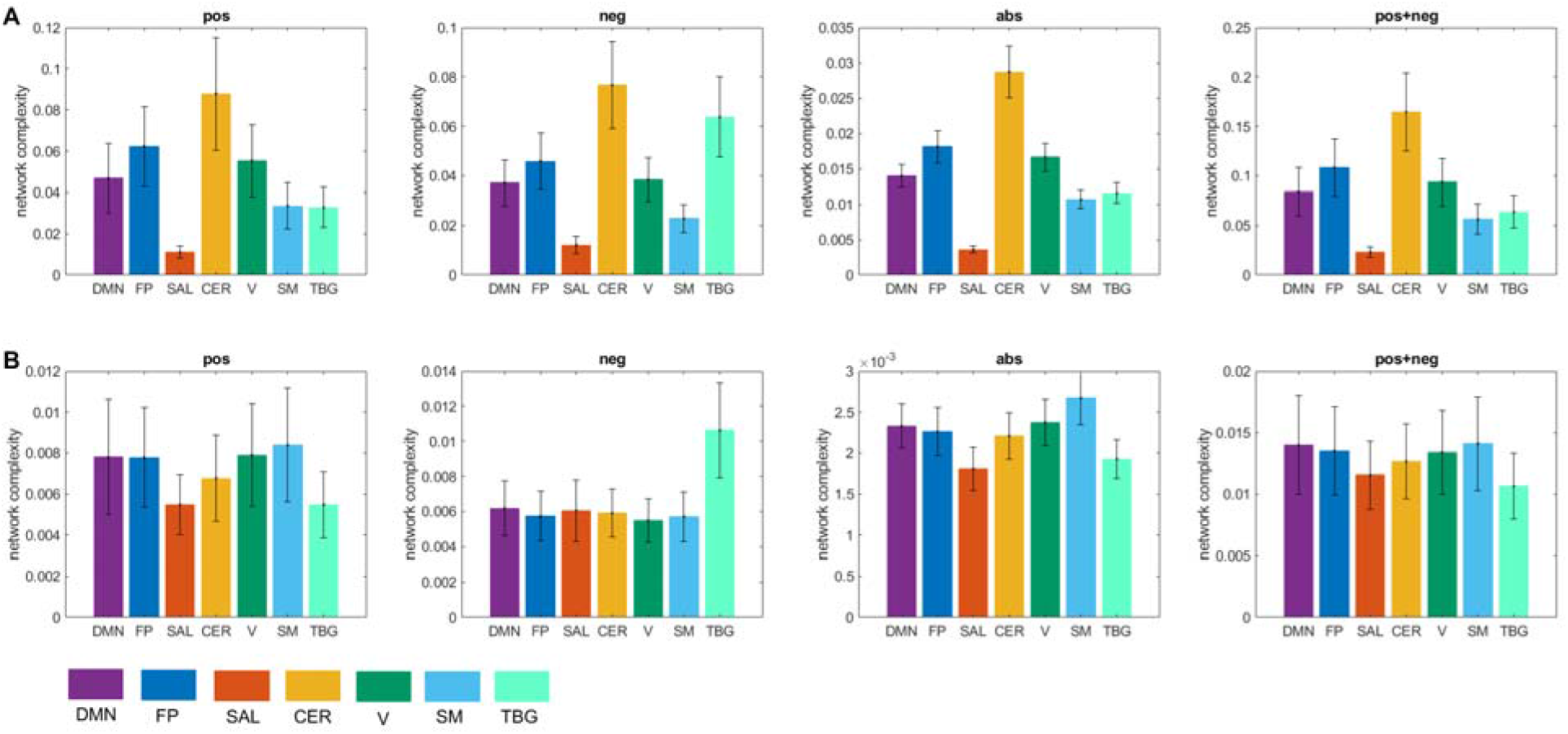
Network complexity. A) Global complexity of seven resting-state networks. B) Global complexity divided by the number of areas in each network. Seven resting-state networks (see Supplementary Table 1) are represented through different colors: default mode network (DMN), frontoparietal (FP), salience (SAL), sensorimotor (SM), visual (V), cerebellar (CER), and temporo-basal-ganglial (TBG) networks. The gray color represents areas not assigned (NA) to any of these networks.

Along these lines, hemispherical differences can be investigated as well. Previous studies have found interhemispheric asymmetry in brain connectivity during resting-state (Medvedev, 2014). We found that the left hemisphere was significantly more complex than the right hemisphere for the seven resting-state networks (figure 10).

**Figure 10.**
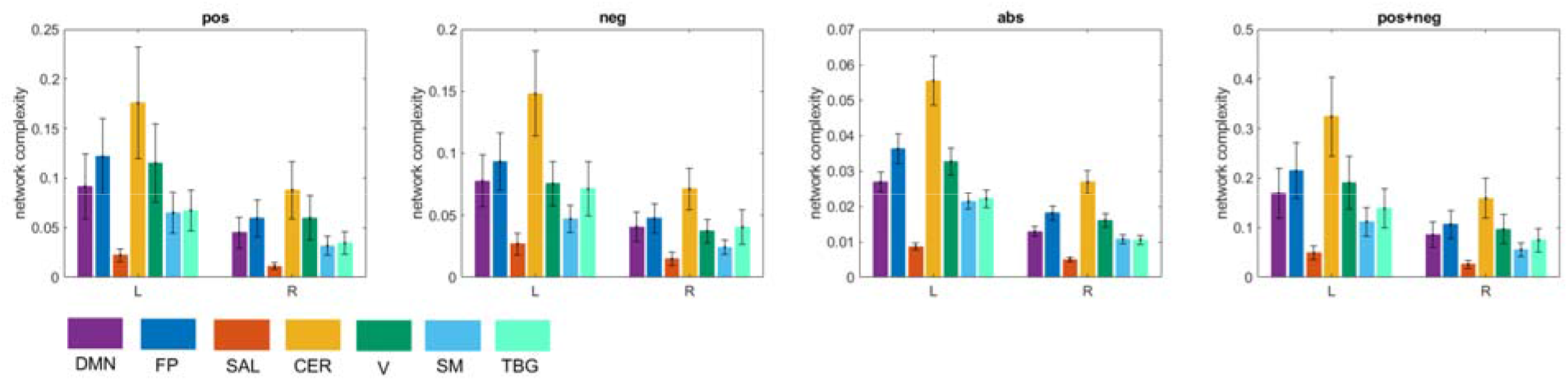
Interhemispheric asymmetry of global complexity. L-left hemisphere, R-right hemisphere. Seven resting-state networks (see Supplementary Table 1) are represented through different colors: default mode network (DMN), frontoparietal (FP), salience (SAL), sensorimotor (SM), visual (V), cerebellar (CER), and temporo-basal-ganglial (TBG) networks. The gray color represents areas not assigned (NA) to any of these networks.

## Discussion

In this study we proposed a new measure of network (global) complexity that is constructed as the sum of the complexities of its nodes. The complexity of each node (i.e, local complexity) was estimated as an index that compares the sample entropy of the time series generated by the movement of a random walker on the network resulting from removing the node and its connections, to the sample entropy of the time series obtained from a regular lattice (the ordered state) and an Erdos-renyi network (disordered state). Our simulations demonstrated that our measure of complexity (equations (7)–(8)), achieves a minimum for the regular lattice and Erdos-Renyi networks, and a maximum at some intermediate state, representing a small-world network with both order and disorder characteristics (figure 2). The rationale behind the use of random walks is that diffusion process are capable of uncovering the large-scale topological structure of complex networks (Noh & Rieger, 2004; Simonsen, Astrup Eriksen, Maslov, & Sneppen, 2004; Skardal & Adhikari, 2018). For instance, random walks are the basis of Infomap (Rosvall & Bergstrom, 2008), a popular method for detecting community structure in complex networks. Past studies of anatomical and functional brain connectivity have found interlinked communities that form a partly decomposable modular architecture (Ashourvan, Telesford, Verstynen, Vettel, & Bassett, 2019; Meunier, Lambiotte, Fornito, Ersche, & Bullmore, 2009). Such architectures are hallmarks of complex systems and are thought to be of fundamental importance for understanding mental processing and cognition (Bola & Borchardt, 2016). In the brain, hierarchies of linked communities span across several levels including brain regions, functional circuits and large-scale networks. This structural diversity cannot be captured by previous structural complexity measures relying mainly on Shannon entropy (Shannon, 1948), but can be probed using random walks (Rosvall & Bergstrom, 2008).

Once we constructed the time series of the random walker’s movement in the network, we needed a measure to estimate its complexity. There is a diversity of complexity measures based on different entropy definitions, such as: Shannon entropy (Shannon, 1948), Tsallis entropy (Tsallis, 1988), spectral entropy (Inouye et al., 1991), wavelet entropy (Rosso et al., 2001), approximate entropy (Pincus, 1991), sample entropy (Richman & Moorman, 2000), fuzzy entropy (Weiting Chen, Zhizhong Wang, Hongbo Xie, & Wangxin Yu, 2007), and permutation entropy (Bandt & Pompe, 2002). In this work we selected sample entropy since it can quantify the amount of regularity and unpredictability of fluctuations in a time series (Richman & Moorman, 2000). This is important because of the presence of communities in brain networks (Ashourvan et al., 2019; Meunier et al., 2009), which will result in repetitive patterns of nodes in the time series of the random walker’s movement (Fortunato & Hric, 2016).

Our study of brain complexity found interhemispheric asymmetry, where the left hemisphere was significantly more complex than the right hemisphere, for all the seven brain networks explored. Previous studies have also found interhemispheric asymmetry in brain connectivity during resting-state. For instance, a recent study used near-infrared spectroscopy (NIRS) signals to estimated functional connectivity matrices (Medvedev, 2014). Their results revealed significantly stronger and denser connectivity patterns in the right hemisphere in most subjects. This denser pattern of connections in the right hemisphere compared to the left hemisphere can lead to a lower structural complexity if it is not accompanied with a significant increase in the entropy of the network. Thus, the balance between the entropy of the network and its density determines the network’s complexity. This was exemplified in figure 3 where we found that the entropy of the positive network and the anti-correlated network were essentially the same, but the positive network was sparser, which resulted in it being more complex than the anti-correlated network.

## Acknowledgments

This work was partially supported by grant RGPIN-2015-05966 from Natural Sciences and Engineering Research Council of Canada. Data were provided [in part] by the Human Connectome Project, WU-Minn Consortium (Principal Investigators: David Van Essen and Kamil Ugurbil; 1U54MH091657) funded by the 16 NIH Institutes and Centers that support the NIH Blueprint for Neuroscience Research; and by the McDonnell Center for Systems Neuroscience at Washington University.

## Supplementary Material

**Table S1.** Resting-state networks

| Network | Areas |
| --- | --- |
| Default mode network (DMN) | Frontal_Mid_Orb_L, Frontal_Mid_Orb_R<br>Frontal_Sup_Medial_L, Frontal_Sup_Medial_R<br>Frontal_Med_Orb_L, Frontal_Med_Orb_R<br>Cingulum_Post_L, Cingulum_Post_R<br>Angular_L, Angular_R<br>Precuneus_L, Precuneus_R |
| Frontoparietal (FP) | Frontal_Sup_L, Frontal_Sup_R<br>Frontal_Mid_L, Frontal_Mid_R<br>Frontal_Inf_Oper_L, Frontal_Inf_Oper_R<br>Frontal_Inf_Tri_L, Frontal_Inf_Tri_R<br>Frontal_Inf_Orb_L, Frontal_Inf_Orb_R<br>Parietal_Sup_L, Parietal_Sup_R<br>Parietal_Inf_L, Parietal_Inf_R<br>SupraMarginal_L, SupraMarginal_R |
| Salience (SAL) | Insula_L, Insula_R<br>Cingulum_Ant_L, Cingulum_Ant_R |
| Sensorimotor (SM) | Precentral_L, Precentral_R<br>Rolandic_Oper_L, Rolandic_Oper_R<br>Postcentral_L, Postcentral_R<br>Supp_Motor_Area_L, Supp_Motor_Area_R |
| Visual (V) | Calcarine_L, Calcarine_R<br>Cuneus_L, Cuneus_R<br>Lingual_L, Lingual_R<br>Occipital_Sup_L, Occipital_Sup_R<br>Occipital_Mid_L, Occipital_Mid_R<br>Occipital_Inf_L, Occipital_Inf_R<br>Fusiform_L, Fusiform_R |
| Cerebellar (CER) | Cerebellum_Crus1_L, Cerebellum_Crus1_R<br>Cerebellum_Crus2_L, Cerebellum_Crus2_R<br>Cerebellum_3_L, Cerebellum_3_R<br>Cerebellum_4_5_L, Cerebellum_4_5_R<br>Cerebellum_6_L, Cerebellum_6_R<br>Cerebellum_7b_L, Cerebellum_7b_R<br>Cerebellum_8_L, Cerebellum_8_R<br>Cerebellum_9_L, Cerebellum_9_R<br>Cerebellum_10_L, Cerebellum_10_R<br>Vermis_1_2, Vermis_3, Vermis_4_5, Vermis_6<br>Vermis_7, Vermis_8, Vermis_9, Vermis_10 |
| Temporo-basal-ganglial (TBG) | Hippocampus_L, Hippocampus_R<br>ParaHippocampal_L, ParaHippocampal_R<br>Amygdala_L, Amygdala_R,<br>Caudate_L, Caudate_R<br>Putamen_L, Putamen_R<br>Pallidum_L, Pallidum_R |

